# GDC: An Integrated Resource to Explore the Pathogenesis of Hearing Loss through Genetics and Genomics

**DOI:** 10.1101/2024.08.19.608726

**Authors:** Hui Cheng, Xuegang Wang, Mingjun Zhong, Jia Geng, Wenjian Li, Kanglu Pei, Yu Lu, Jing Cheng, Fengxiao Bu, Huijun Yuan

## Abstract

Effective research and clinical application in audiology and hearing loss (HL) often require the integration of diverse data. However, the absence of a dedicated database impeded understanding and insight extraction in HL. To address this, the Genetic Deafness Commons (GDC) was developed by consolidating extensive genetic and genomic data from 51 public databases and the Chinese Deafness Genetics Consortium, encompassing 5,983,613 variants across 201 HL genes. This comprehensive dataset detailed the genetic landscape of HL, identifying six novel mutational hotspots within DNA binding domains of transcription factor genes, which were eligible for evidence-based variant pathogenicity classification. Comparative phenotypic analyses highlighted considerable disparities between human and mouse models, with only 130 human HL genes exhibiting hearing abnormality in mice. Moreover, gene expression analyses in the cochleae of mice and rhesus macaques demonstrated a notable correlation (R^2^ = 0.76). Utilizing gene expression, function, pathway, and phenotype data, a SMOTE-Random Forest model identified 18 candidate HL genes, including *TBX2* and *ERCC2*, newly confirmed as HL genes. The GDC, as a comprehensive and unified repository, significantly advances audiology research and clinical practice by enhancing data accessibility and usability, thereby facilitating deeper insights into hearing disorders.

## Introduction

Hearing loss (HL) is the most common sensory deficit and one of the most common congenital abnormalities. Estimates indicate that among every 1,000 newborns screened, 1.1 to 3.5 will have HL ^1,2^. The etiology of HL is multifactorial, encompassing genetic defects, physical trauma, infections, drug toxicity, noise exposure, and aging, among other factors ^3,4^. Genetic predispositions play a pivotal role in congenital and early-onset HL, characterized by significant genetic and phenotypic heterogeneity, with more than 200 genes identified thus far ^5–7^.

Current research and clinical applications in audiology and hearing disorders, such as novel HL gene identification, auditory mechanism exploration, research on the auditory system development, variant interpretation and genetic diagnosis, and the development of gene therapies, hold the promise of translating individual genomic data into clinically relevant information to aid in disease diagnostics and facilitate personalized therapeutic decision making ^8–12^. These efforts depend critically on the integration of vast data and knowledge accumulated across numerous databases and repositories. This integration includes gene/region-level annotations detailing genomic features (e.g., UCSC genome browsers ^13^ and ENSEMBL ^14^), transcriptional information (e.g., gEAR ^15^ and GTEx ^16^), gene/protein functions (e.g., UniProtKB ^17^ and Gene Ontology ^18^) and structures (e.g., InterPro ^19^ and PDB ^20^), pathway information (e.g., Reactome ^21^ and KEGG ^22^), gene-gene interactions (e.g., STRING ^23^ and BioGrid ^24^), and gene-drug interactions (e.g., PharmGKB ^25^ and DGIdb ^26^). Variant annotations also play a pivotal role in interpreting the effects of molecular processes and disease causation, including population frequencies (e.g., gnomAD ^27^, ChinaMAP ^28^, and BBJ ^29^), genotype-phenotype correlations and pathogenicity classifications (e.g., ClinVar ^30^, HGMD ^31^, and DVD ^5^), in silico function predictions (e.g., CADD ^32^, REVEL ^33^, and SpliceAI ^34^), and automated literature mining (e.g., LitVar ^35^ and Variant2Literature ^36^). Additionally, disease and phenotype information extracted from literature through expert curation includes standardized disease/phenotype descriptions (e.g., HPO ^37^ and MedlinePlus ^38^), disease-gene correlation (e.g., OMIM ^39^ and ClinGen ^40^), and animal models (e.g., MGI ^41^). Together, these diverse databases provide essential support for researchers to explore the complexities of genetic and molecular mechanisms underlying audiology and hearing disorders, offering crucial insights into broader biological processes, disease pathogenesis, and potential therapeutic interventions.

However, the lack of a dedicated, comprehensive database for hearing research has significantly hindered the compilation and integration of these resources. Existing hearing-related databases like DVD and gEAR cater to niche aspects of hearing loss research, with DVD focusing on variant classification within HL genes and gEAR on gene expression in the cochlea. Information is often scattered across specialized catalogs tailored to specific fields, particular model organisms, or specific techniques, resulting in heterogeneous data vocabularies, ontologies, and formats. It complicates efforts to fully reconcile and integrate the data, posing significant challenges in understanding the landscape of hearing loss, identifying and prioritizing relevant information, and consequently impeding the extraction of meaningful insights.

To bridge the gap, we established Genetic Deafness Commons (GDC, http://gdcdb.net/), a standardized database and knowledge base that comprehensively consolidates and integrates genetic and genomic data from both public and in-house sources. The GDC leveraged over 51 public databases to provide extensive information on HL genes, variants, and phenotypes. Additionally, it integrates genetic findings from 22,125 HL cases from the Chinese Deafness Genetics Consortium (CDGC), offering real-world patient cohort data to support variant interpretation and curation. Utilizing the extensive dataset of GDC, this study conducts a thorough analysis of the genetic architecture of HL genes and variants, uncovering multiple new patterns to improve the efficacy of variant pathogenicity interpretation and genetic diagnosis. By integrating public gene expression data from mice and in-house data from rhesus macaques, this study applied the machine learning method to identify a set of new candidate HL genes, demonstrating the significant value of GDC in deciphering the genetic and molecular foundations of audiology and hearing-related conditions.

## Results

### Overview of GDC

The current version of the GDC amalgamated data from 51 public sources and the CDGC project, incorporating 5,983,613 variants across 201 reported HL genes (Figure 1). GDC encompasses detailed disease information, gene expression patterns, annotations related to gene functions and pathways, protein structures and domains, gene-gene and gene-drug interactions, as well as other essential data. Additionally, GDC introduces multiple specialized annotations to enhance understanding of each variant, including allele frequencies from 14 populations of five large genome sequencing studies, variant pathogenicity classifications from four sources, 20 prediction scores for functional damage, genotype-phenotype correlations, and variant-based literature mining results. Furthermore, GDC offers a user-friendly query interface to ensure that information is readily accessible in various formats such as graphical representations, tables, and downloadable files, facilitating easy search and navigation for researchers investigating any HL-related variants and genes.

**Figure 1.**
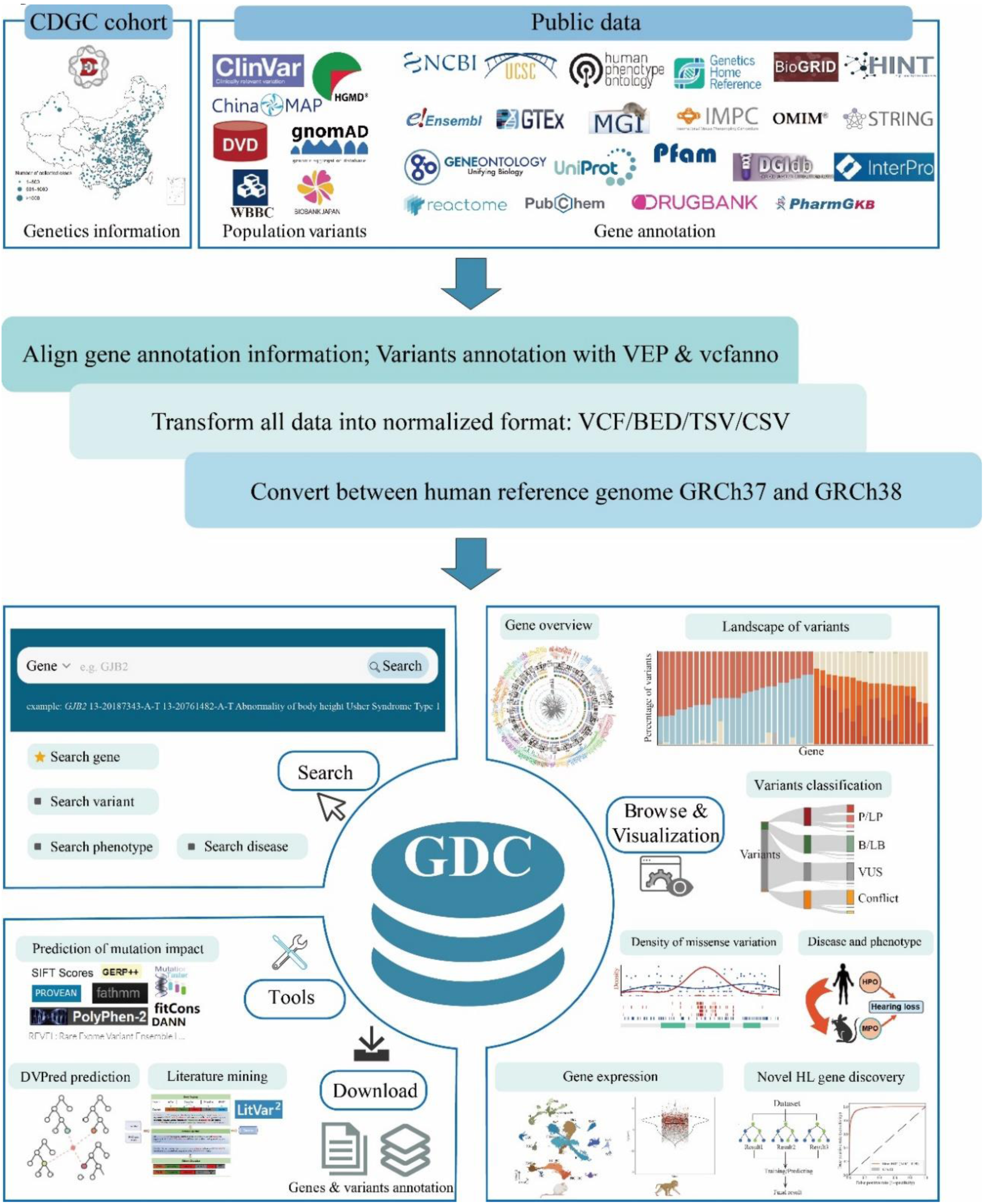
Construction of the Genetic Deafness Commons (GDC). The data from 51 public databases and the Chinese Deafness Genetics Consortium were collected. A series of data processing, including data alignment, gene and variant annotation, format transformation, and lifting over between human reference genome GRCh37 and GRCh38, was applied to integrate and harmonize data in GDC.

Of the 201 HL genes in the GDC, 139 were associated with non-syndromic HL (NSHL) and 93 with syndromic HL (SHL), linking to 92 different disorders including Usher syndrome, Alport syndrome, and Waardenburg syndrome (Figure 2A). Notably, 31 genes were implicated in both NSHL and SHL. In terms of inheritance patterns, 95 genes were linked to autosomal dominant (AD), 128 to autosomal recessive (AR), eight to X-linked (XL), and one to mitochondrial (MT) inheritance patterns. Furthermore, 31 genes exhibit both AD and AR inheritance patterns. These genes are annotated with 7,656 functional terms (with a median of 27 terms per gene) and 8,695 phenotypic terms (with a median of 19 terms per gene).

**Figure 2.**
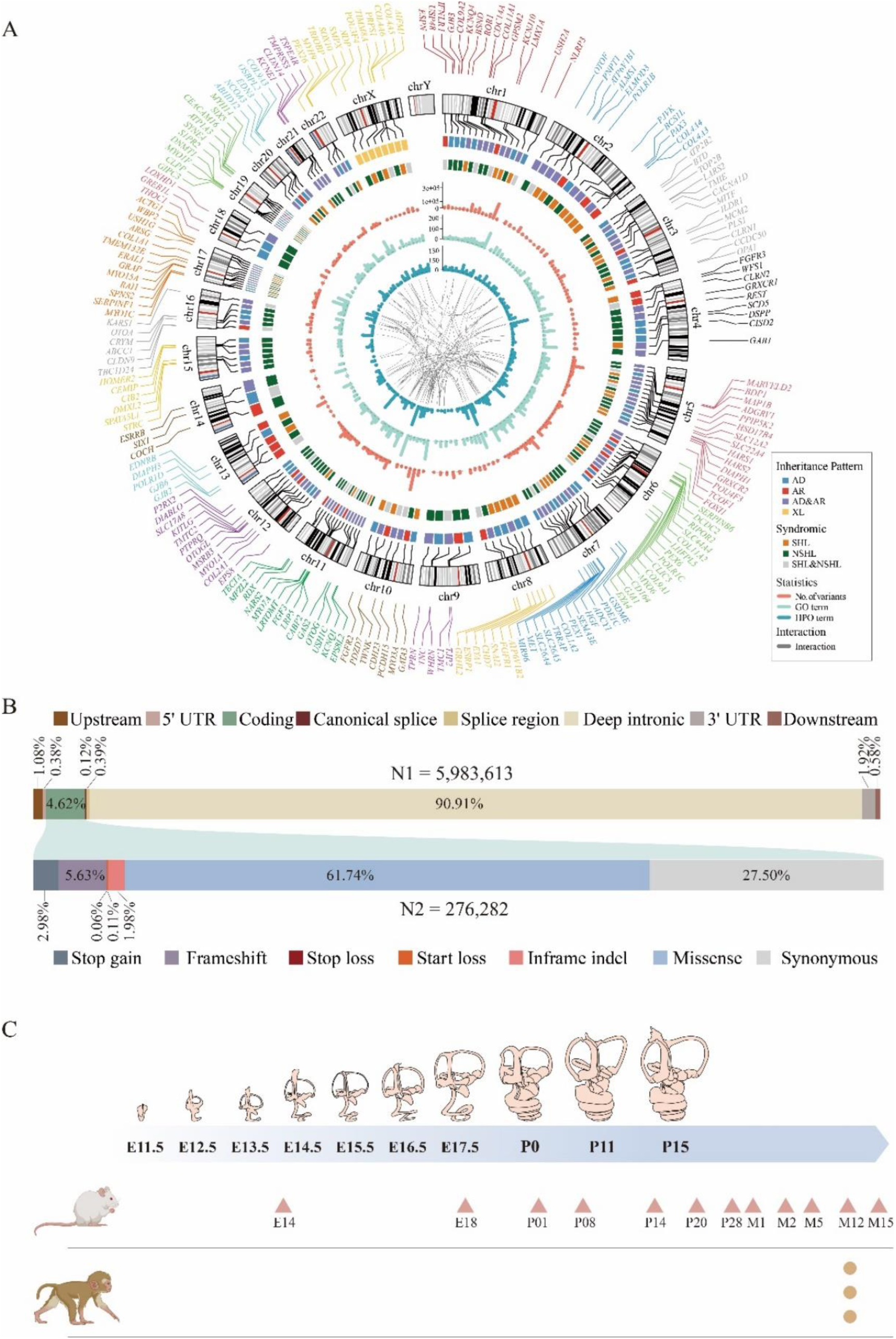
Summary of genetic and genomic information in GDC. (A) Summarization of 201 HL genes. The outer circle is genomic location of 201 genes. Genes located on different chromosomes are distinguished by color. Inner circles were inheritance pattern, syndromic HL or not, statistics of variants, GO term, and HPO term, and gene-gene interactions, respectively. (B) The total number and proportion of variants observed classified by genomic location. Variants in coding regions were further classified by functional consequence. (C) Developmental time points of gene expression data from cochleae of mice and rhesus macaques that integrated by GDC.

Within the exons, introns, and 1kb upstream and downstream regions of the 201 HL genes, 5,983,613 genetic variations were collated in the GDC from multiple sources including CDGC (v202407, n = 217,915), gnomAD (v3.1, n = 4,973,216), ChinaMAP (v1, n = 997,781), DVD (v9, n = 2,100,721), ClinVar (20230702, n = 121,439), and HGMD (2023v2, n = 24,470). Of all variants, 4.63% were located in exonic and adjacent (±8bp) intronic regions. Missense variants constitute the majority of such variants at 61.74%. The next most prevalent types are synonymous variants (27.5%) followed by indels (5.63% frameshift and 1.98% in-frame), stop gain (2.98%), 5’ UTR (0.38%), canonical splice sites (±2 bp of an intron, 0.12%), splice regions (±3-8 bp of an intron, 0.39%), 3’ UTR (1.92%), up/downstream regions (1.68%), and start/stop loss (<0.2%) (Figure 2B). Most variants were extremely rare, with the GDC dataset containing 1,436,357 (24%) variants not reported in gnomAD. Additionally, 2,338,525 (39.06%) variants in the GDC dataset are singletons in gnomAD, with doubletons, tripletons, and quadruplets accounting for 10.72%, 5.15%, and 3.08% respectively, and 639,113 variants (10.67%) have a minor allele frequency (MAF) < 0.0002 and allele count (AC) > 4.

Furthermore, GDC incorporated extensive gene expression data from both human and model animals. This included bulk RNA-seq data of 54 human tissues sourced from the GTEx database, single-cell RNA-seq data at 12 developmental stages of the mouse cochlea from five public repositories (GSE172110, GSE182202, GSE181454, GSE202920, and CRA004814) ^42–46^, and in-house bulk RNA-seq data from three adult rhesus macaque cochlea (Figure 2C). These datasets enable the GDC to provide a detailed and dynamic view of gene expression across different species and developmental stages, significantly enhancing the potential for discoveries in audiology and hearing disorders.

### Variant Pathogenicity Classification and Curation

Across the CDGC, DVD, ClinVar, and HGMD databases, 38.18% (n = 2,284,692) of all GDC variants were classified as pathogenic (P), likely pathogenic (LP), benign (B), likely benign (LB), or variant with uncertain significance (VUS). A total of 35,702 variants were reported as P/LP by at least one source (Figure 3A). Among these variants, 16,244 (45.5%) were consistently classified as P/LP across two or more sources, whereas 11,607 (32.51%) P/LP variants were reported in only one source. Conversely, 7,851 (21.99%) variants showed medically significant classification conflicts, shifting between P/LP and B/LB, or VUS, indicating the need for further validation using more extensive patient data (Figure 3B). Among the databases, HGMD exhibited the highest incidence of 1,040 conflicts between P/LP and B/LB classifications, as shown in Figure 3C. Remarkably, 640 previously classified as ‘P/LP’ (‘DM’ or ‘DM?’) variants in HGMD were reclassified to B/LB by CDGC.

**Figure 3.**
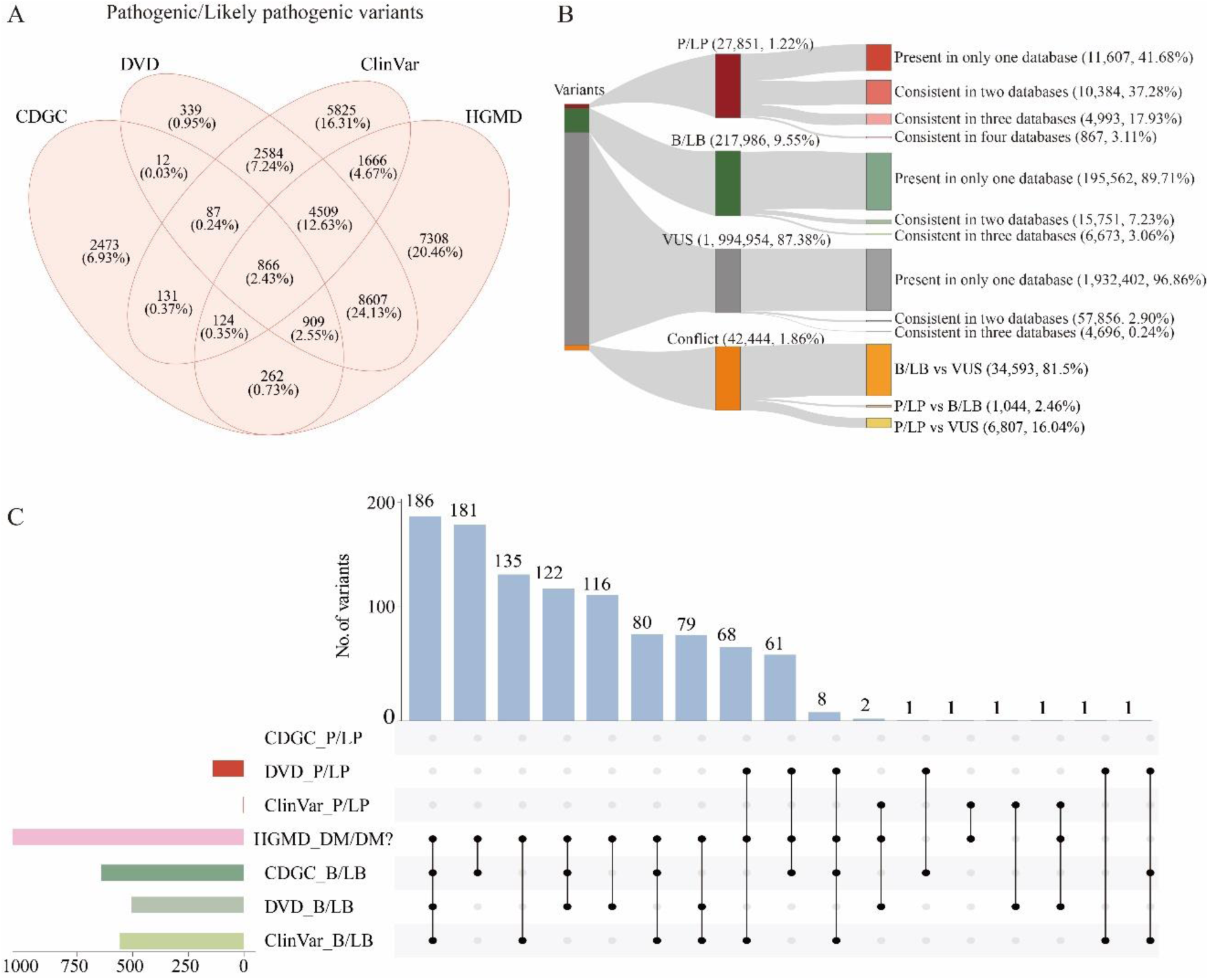
Variant pathogenicity classification from varied sources. (A) Shared and unique P/LP variants collected from DVD, ClinVar, HGMD and CDGC datasets. (B) Comparative overview of variants classification in GDC and the consistency among the databases. (C) P/LP and B/LB conflictions across DVD, ClinVar, HGMD and CDGC. Bar plot on left side shows total number of conflicted variant classifications for each dataset.

### Analyzing Pathogenic Variants for Insights into Gene Function and Tolerance

Protein truncating variants (PTVs), including stop-gain, frameshift, start loss, stop loss, and canonical splice site changes, are expected to result in a complete loss of function (LoF) of the affected transcripts and are considered potentially deleterious ^47^. Within the protein-coding genes included in the GDC, the count of PTVs ranged from 6 to 1,886. A total of 31,558 PTVs were classified by CDGC, DVD, ClinVar, or HGMD. Notably, the majority of the PTVs classified as VUS (n =6,785) were contributed by DVD (77.47%). Analysis of the variant types among P/LP classifications for each gene revealed that PTVs constitute more than half of P/LP variants in 127 genes (67.91%) (Figure 4A). All P/LP variants in the gene *EPS8* (n=13), *SYNE4* (n=12), *MPZL2* (n=11), *GRXCR2* (n=5), *BDP1* (n=3), *GAS2* (n=2), *CLDN9* (n=1), and *CD164* (n=1) were PTVs, indicating that LoF is likely the disease mechanism in these genes. In 52 genes (27.96%), more than half of the P/LP variants were missense (Figure 4B). Specifically, in genes like *DIABLO*, *ELMOD3*, *IFNLR1*, *MYO1C*, *P2RX2*, *PDE1C*, *PLS1*, *POLR1B*, *S1PR2*, *SCD5*, *THOC1*, and *WBP2*, all P/LP variants were missense. By literature review, a gain of function (GoF) mechanism was only reported in *DIABLO* ^48^, suggesting further exploration of GoF as the probable disease mechanism across these genes. By analyzing the rates of P/LP variants among PTVs for each gene, the distribution of genes across different inheritance patterns is visualized. Conversely, genes tolerant to LoF, such as *DIPAH3* and *COCH*, are found in a quadrant where both ratios are below 50%. It is noteworthy that genes with potential lethal LoF variants, like *ACTG1* and *AIFM1*, also fall into this category.

**Figure 4.**
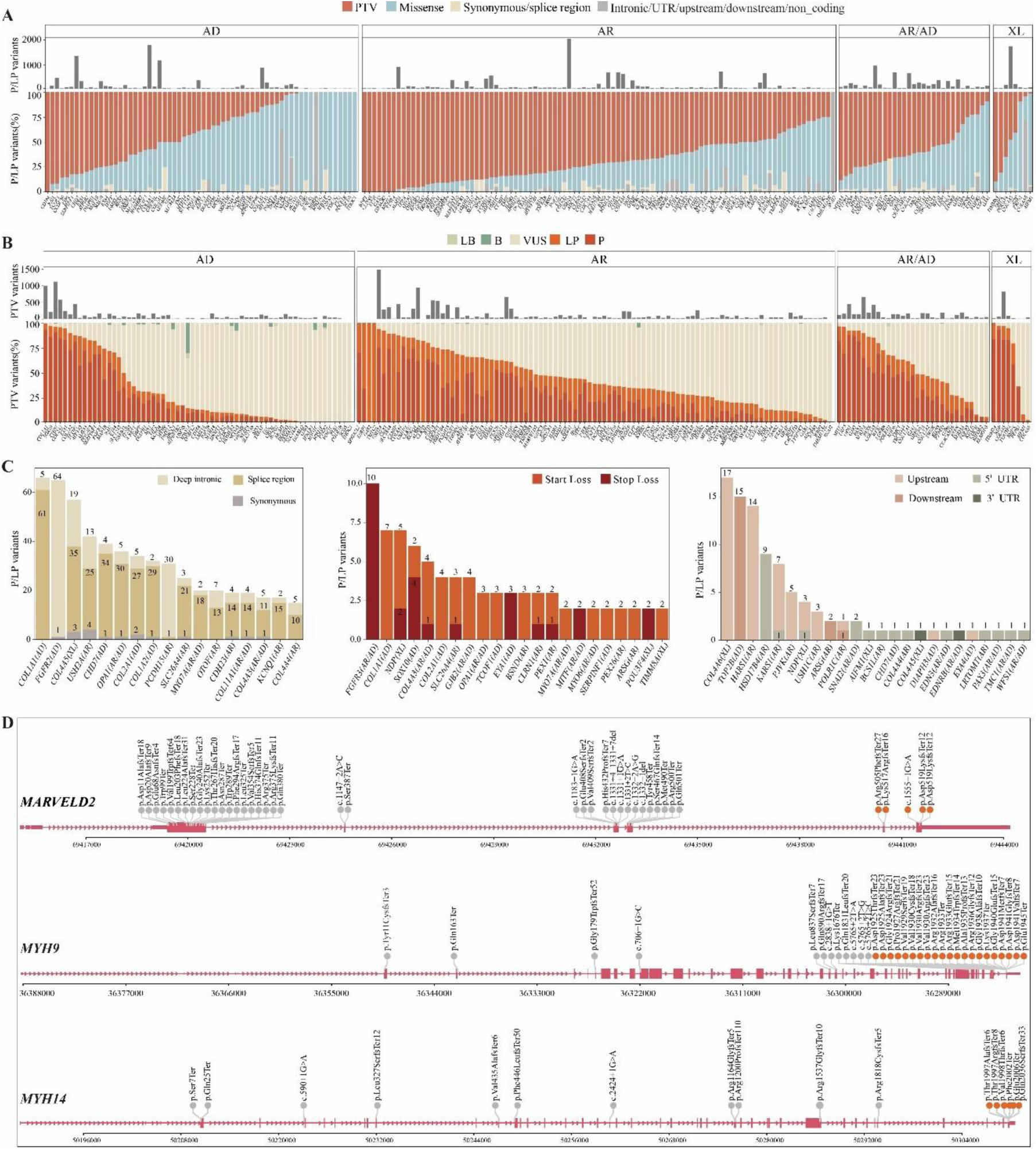
Genetic landscape of HL-associated genes. (A) Variant types of P/LP variants across 201 HL genes. The barplot above indicates the number of variants. Abbreviations: AD, Autosomal dominant; AR, Autosomal recessive; XL, X-linked; PTV, Protein Truncating Variant. (B) Pathogenicity classification of PTVs across 201 HL genes. (C) Distribution of rare types of pathogenic variants, including deep intronic, synonymous, splice region, start loss, stop loss, UTRs and intergenic. (D) Distribution of PTVs in genes enriched with NMD-escape PTVs

In addition, rare types of pathogenic variants were observed in multiple genes (Figure 4C). For instance, pathogenic splicing variants are prevalent in genes such as *COL1A1*, *FGFR2*, *COL4A5*, and *USH2A*. Loss-of-start and loss-of-stop variants are typically found in genes *FGFR3*, *COL1A1*, and *NDP*. Furthermore, pathogenic variants affecting upstream regions, untranslated regions (UTRs), or downstream non-coding regions are observed in genes *COL4A6*, *TOP2B*, and *HARS1*.

In certain genes, *MARVELD2*, *MYH9*, *MYH14*, *CABP2*, *CLIC5*, *HOMER2*, *SPATA5L1*, and *WFS1*, pathogenic PTVs were abundantly identified within the last exon or the final 50 base pairs of the penultimate exon (Figure 4D), regions predicted to potentially escape nonsense-mediated mRNA decay (NMD). The pathogenicity of 3’-end PTVs in these genes requires careful evaluation, given their potentially significant impact on protein activity and stability.

### Identification of ‘Hotspots’ for Pathogenic Missense Variants

We further evaluate the distribution of missense P/LP variants using Gaussian kernel density estimation, overlapping with protein domains to identify hotspots of disease-related missense mutations. By integrating various sources, we identified 8,787 missense P/LP variants in 201 HL genes (ranging from 1 to 785 per gene). Significant enrichment of P/LP missense variants was predominantly found in transcription factor genes, including *GATA3* (NP_002042: p.264-342), *GRHL2* (NP_079191: p.213-438), *LMX1A* (NP_796372: p.196-252), *MITF* (NP_000239: p.198-288), *PAX3* (NP_852124: p.34-159; p.220-276), *POU3F4* (NP_000298: p.186-340), *POU4F3* (NP_002691: p.179-333), *REST* (NP_001350382: p.159-412), *SIX1* (NP_005973: p.127-181), and *SOX10* (NP_008872: p.103-172).

Remarkably, these enriched regions were all DNA-binding domains (Figure 5A). The exceptions in transcription factor genes were *FOXI1*, *SIX5*, and *SNAI2*, likely due to the limited number of identified P/LP variants. We calculated the positive likelihood ratio for a missense variant being P/LP in these hotspot regions. The lower boundary of the 95% confidence interval of the positive likelihood ratio (LR+_LB) ranged from 0.26 to 9.75 (Figure 5B). Compared to the thresholds for pathogenic evidence as suggested by Tavtigian et al ^49^, hotspots in *PAX3, SOX10,* and *GATA3* met the moderate level (LR+_LB > 4.3) of the ACMG/AMP pathogenic evidence. *LMX1A*, *POU3F4,* and *MITF* fulfill the supporting level (LR+_LB > 2.08) for pathogenicity according to the same criteria. Additionally, missense variants of *KCNQ1* and *KCNQ4* were significantly enriched in the ion transport domain (LR+_LB = 2.08) and P-loop domain (LR+_LB = 9.78), respectively, consistent with previous reports ^50^. No significant missense variant enrichment was observed in other genes.

**Figure 5.**
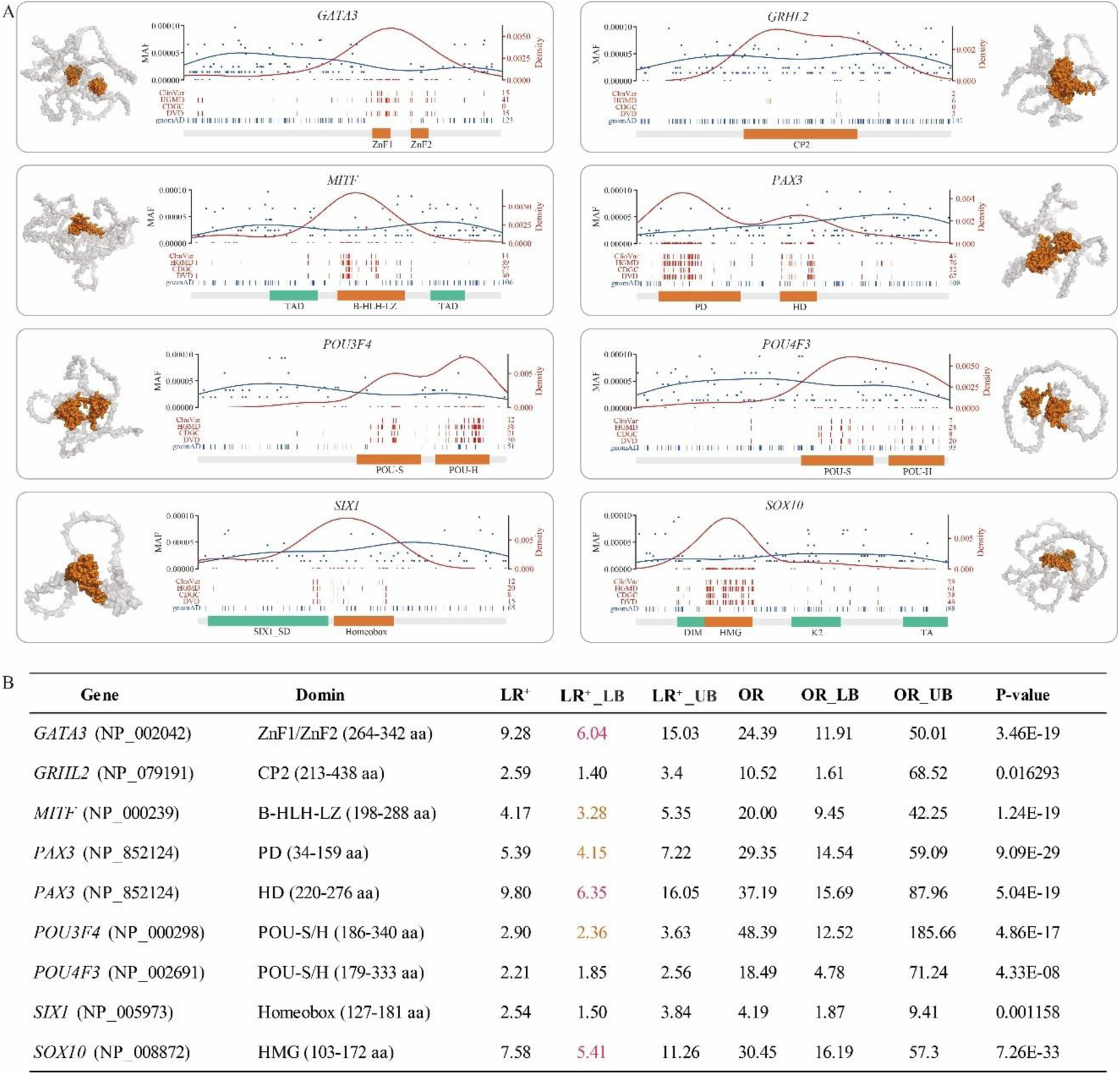
Identification of gene domains for enrichment of pathogenic missense variants. (A) Distribution of P/LP missense variants across genes. Red curves are the density for P/LP variants reported in ClinVar, HGMD, DVD, and CDGC cohort. Blue curves are for variants identified in the gnomAD population. The horizontal bar represents protein domains, with the orange color indicating DNA-binding domains. Abbreviations: ZnF, Zinc finger; CP2, Grh/CP2 DNA-binding domain profile; TAD, Transactivation Domin; bHLH-LZ, Basic helix-loop-helix, Leucine-zipper; PD, ‘Paired box’ domain; HD, Homeodomain; POU-S, POU-specific domain; POU-H, POU-homeodomain; SIX1_SD, Transcriptional regulator, SIX1, N-terminal SD domain; DIM, Dimerization; HMG, High mobility group box; K2, Context-dependent transactivation domain; TA, Transactivation. (B) Statistical analysis of likelihood ratio (LR) and odds ratio (OR) for a missense variant being P/LP in these domains.

### Discrepancies of HL Phenotype between Human and Mouse Models

Phenotypic analysis utilizing model organisms, predominantly mice, is crucial for identifying disease genes and elucidating underlying mechanisms. Nevertheless, phenotypes associated with homologous genes often exhibit substantial differences between humans and mice ^51–53^. Among the 201 human HL genes in GDC, only 130 exhibited an HL phenotype in mice, while 34 were associated with normal hearing (NH), and 37 lacked relevant HL data (NA) (Figure 6A). Notably, mouse models showed greater alignment for genes associated with AR inheritance, while discrepancies in HL phenotypes were noted in 23 AR, 33 AD, 4 XL, and 9 AR/AD genes. We introduced a phenotype similarity (PS) score to systematically quantify and compare phenotypes between humans and mice ^52^. No correlation was observed between the PS score and protein sequence homology (Figure 6B).

**Figure 6.**
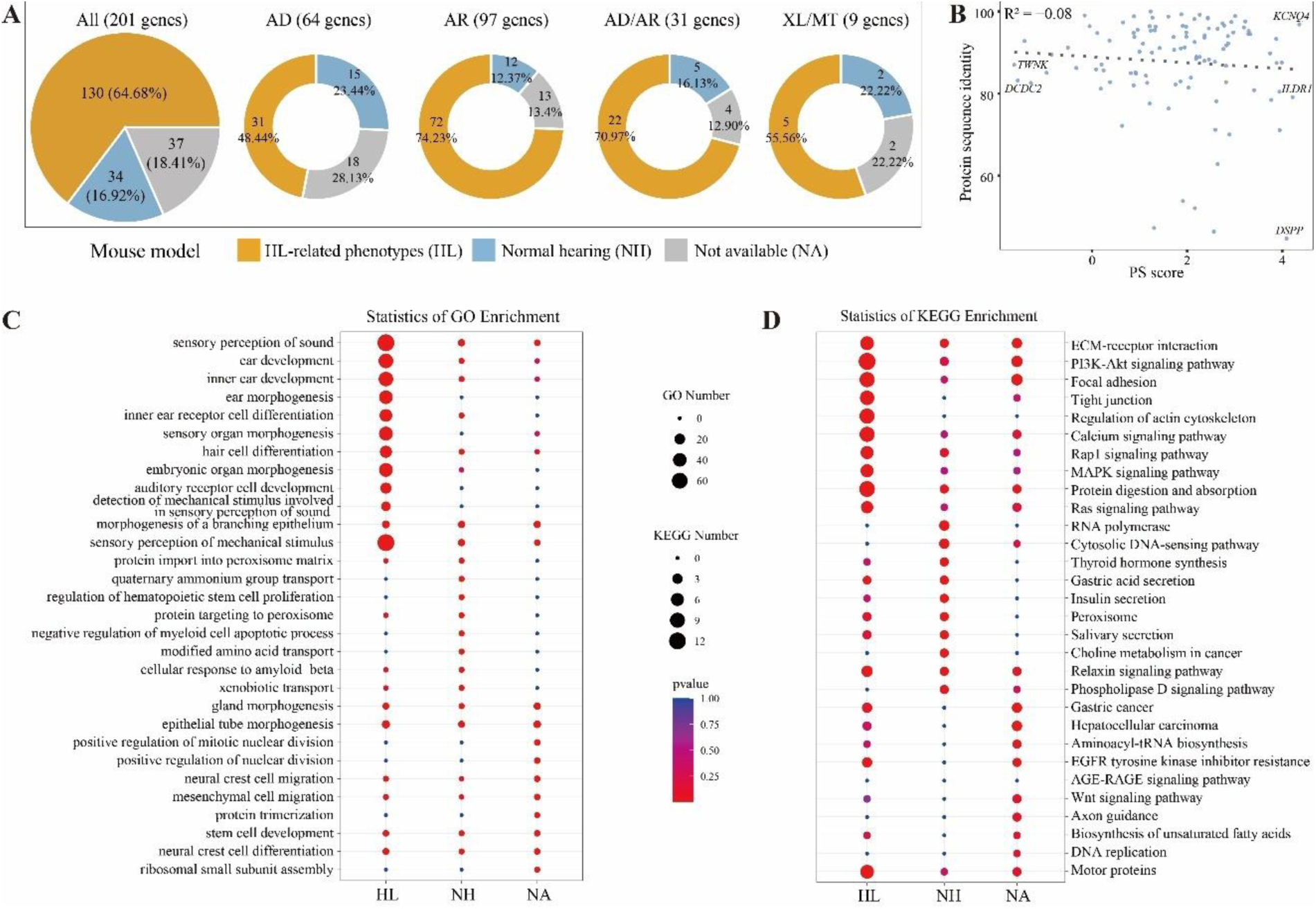
Comparison of human and mouse HL phenotypes. (A) Assessment the hearing phenotypes of 201 human HL genes in mouse models. Abbreviations: HL: hearing loss genes; NH: normal hearing genes; NA: lacked relevant HL data genes. (B) No correlation is observed between the phenotypic similarity score and the protein sequence homology. (C) GO enrichment analysis on HL, NH and NA gene groups. Top 10 enriched GO terms for each group were selected for display. (D) KEGG pathway enrichment analysis on HL, NH and NA gene groups. Top 10 enriched pathways for each group were selected for display.

Further, we explored the differences in gene ontology (GO) and pathways (KEGG) enrichment between genes with consistent and inconsistent HL phenotypes in humans and mice. The HL group demonstrated significant enrichment in GO terms related to sensory perception of sound, ear development, and inner ear receptor cell differentiation (Figure 6C). Conversely, the NH group showed enrichment in functions such as quaternary ammonium group transport and protein targeting to peroxisome, while the NA group primarily displayed enrichment in cell development and migration (Figure 6C) Pathway analysis revealed that the HL group was significantly enriched in the CM-receptor interaction, PI3K-Akt signaling pathway, and MAPK signaling pathway. In contrast, the NH group was notably enriched in the Ras signaling pathway and cytosolic DNA-sensing pathway, whereas the NA group showed significant enrichment in pathways related to hepatocellular carcinoma and AGE-RAGE signaling (Figure 6D). These variations in function and pathway enrichment likely reflect the divergent roles these genes play in mice, highlighting the varied functions genes may exhibit in different biological contexts.

### Expression Features of HL Genes in Mouse and Rhesus Macaque Cochlea

Through gene expression profile analysis on single-cell RNA-seq data at various developmental stages of the mouse cochlea, we identified 21 major cell types, distributed across 12 groups (Figure 7A, 7B). These groups included cells located in and around the organ of Corti, modiolus, Reissner’s membrane, stria vascularis, and spiral ligament, along with multiple types of immune cells. Overall, 201 HL genes were significantly more expressed in the inner ear cells than all other genes. However, differential analysis revealed that HL genes were seldom among the top five most expressed genes in hair cells, supporting cells, or other cell types. Additionally, bulk RNA-seq data from the rhesus macaque cochlea, provided by the GDC, also indicated a generally higher expression of the 201 genes compared to other genes, and the correlation between data from monkeys and mice was significant (R^2^ = 0.76). We also examined genes that did not exhibit an HL phenotype in mice and found no significant difference in gene expression between rhesus macaque and mouse models.

**Figure 7.**
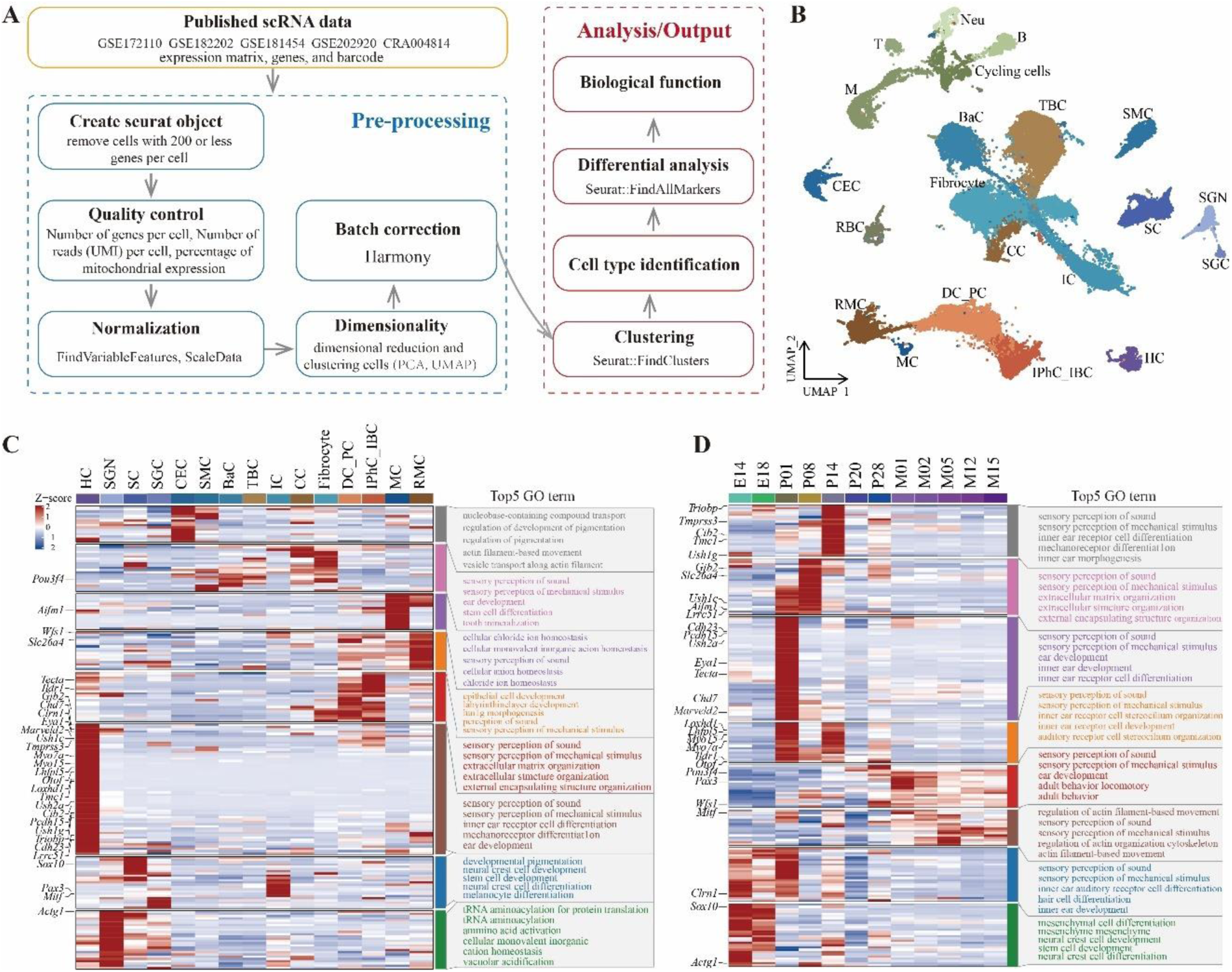
Single-cell transcriptome landscape of mouse cochlea. (A) Analysis process of single cell data of mouse cochlea. (B) The distribution of different cell types in mouse cochlea. Abbreviations: HC, Hair cell; SGN, Spiral ganglion neuron; SGC, Satellite glial cell; SC, Schwann cell; CEC, Capillary endothelial cell; SMC, Smooth muscle cell; MC, Marginal cell; BaC, Basal cell; TBC, Tympanic border cell; IC, Intermediate cell; CC, Chondrocyte; DC_PC, Deiter cell and pillar cell; IPhC_IBC, Inner phalangeal cell/inner border cell; RMC, cells in Reissner’s membrane; T, T cell; B, B cell; M, Macrophage; Neu, Granulocyte/neutrophil; RBC, Red blood cell. (C) The expression distribution of the 201 HL genes in different cell types of the mouse cochlea. (D) The distribution of the 201 HL genes in different developmental stages of the mouse cochlea. Genes were clustered based on expression pattern in (C) and (D). Top five enriched GO terms for each cluster are listed.

Next, we analyzed gene expression across different cell types, clustered 201 HL genes into eight groups, and highlighted the top 30 genes frequently diagnosed in CDGC patients (Figure 7C). Certain genes, including *Cdh23, Cib2, Clrn1, Eya1, Lhfpl5, Loxhd1, Lrrc51, Marveld2, Myo15, Myo7a, Otof, Pcdh15, Tmc1, Tmprss3, Triobp, Ush1c, Ush1g* and *Ush2a*, exhibited high expression levels in hair cells, indicating their crucial roles in inner ear development. *Gjb2, Ildr1*, *and Tecta*, primarily expressed in supporting cells, were vital for maintaining auditory function. Additionally, genes like *Actg1*, *Aifm1*, *Clrn1*, *Pou3f4*, and *Sox10* were expressed in various other cell types, suggesting a broad spectrum of functions. We also examined gene expression across different developmental stages (Figure 7D). *Actg1*, *Clrn1*, and *Sox10* showed high expression during the embryonic stage, indicating their involvement in early inner ear development. Conversely, *Pax3, Pou3f4, Wfs1*, and *Mitf*, showed postnatally high expression, suggesting significant roles in sustaining hearing function during aging.

### Identification of Candidate HL Genes by Integrating Expression and Function Datasets

Utilizing comprehensive expression and functional annotation datasets incorporated by the GDC, we explored novel candidate genes for HL employing a supervised machine learning approach (Figure 8A). Our analysis incorporated the expression levels of genes across 15 cochlear cell types derived from mouse single-cell RNA-seq data and three bulk RNA-seq experiment results on rhesus macaque cochlea. Additionally, we computed the expression correlations among all genes, utilizing the 201 coefficients correlated with HL genes as features. Along with 30 pathway annotations and 30 GO annotations, it constitutes a total of 275 features to be fed into the subsequent model.

**Figure 8.**
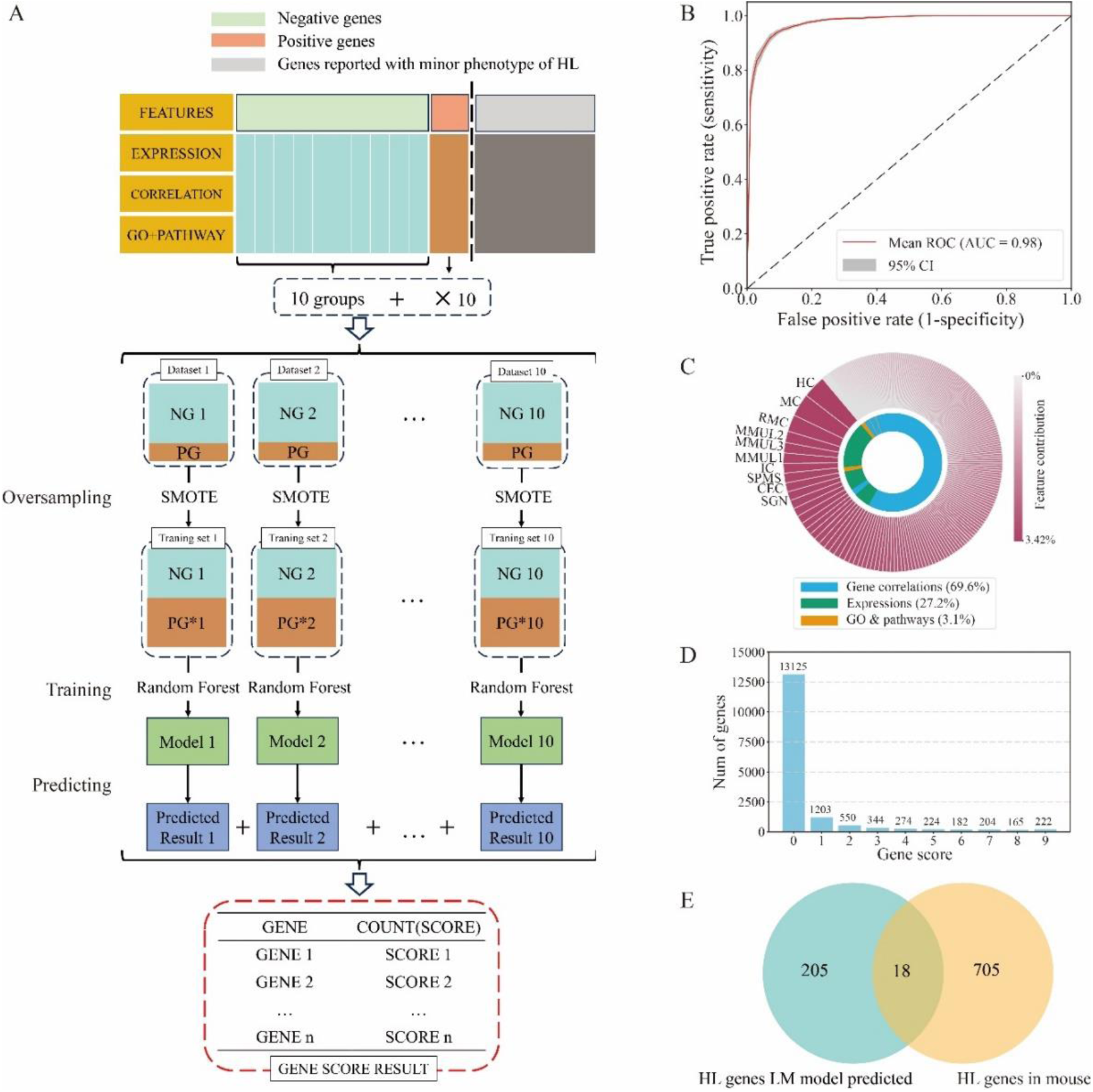
Candidate HL genes discovery via SMOTE Random Forest Model. (A) Workflow for gene classification and prediction model. SMOTE method was used for oversampling to create training sets. Abbreviations: NG, HL-negative genes; PG, HL-positive genes. (B) Mean ROC Curve with 95% Confidence Interval (CI) for 10 SMOTE-Random Forest classifiers. The red line represents the mean ROC curve. The shaded grey region denotes the 95% CI. (C) Feature contribution to the SMOTE-Random Forest model. The outer ring shows the importance of each feature. The top 10 features are labeled with text. The inner ring categorizes the features into three groups: gene correlation, GO & pathways, and expression. (D) Distribution of gene scores. The x-axis represents the gene scores for HL. The y-axis indicates the number of genes in each gene score category. (E) Overlap between genes with a high score of 9 and genes showing HL phenotype in mouse model.

By excluding genes without expression information, the dataset started with 18,771 genes. To construct the training set, we initially selected 201 HL-associated genes for the positive dataset, while the remaining genes constituted the negative dataset. To improve phenotype association accuracy, we excluded four genes from the positive dataset that had been refuted by ClinGen and removed 2,081 genes from the negative dataset that were listed in the Human Phenotype Ontology (HPO) as having HL as a minor symptom. This refinement yielded a dataset of 197 HL-positive genes and 16,493 HL-negative genes. Given the significant data imbalance, we implemented the SMOTE-Random Forest classifier. We randomly divided the 16,493 HL negative genes into ten groups, each paired with the 197 HL positive genes to form ten distinct datasets. We then trained these datasets and evaluated them using a split of 0.2 for the test set and 0.8 for the training set. All ten models demonstrated high performance metrics (Figure 8B), achieving precision scores ranging from 0.90 to 0.95 (mean of 0.93), recall scores from 0.87 to 0.95 (mean of 0.925), and F1 scores from 0.91 to 0.94 (mean of 0.924). The models also provided insights into feature importance. Among the top 10 important features, the majority (9 out of 10) were related to expressions, including gene expression data in six mouse cochlea cell types and three rhesus macaque cochlea samples, with the only one exception being the GO term of sensory perception of mechanical stimulus. Although expression correlations with the known 197 HL genes were not ranked among the top features, they accounted for a significant proportion of the overall feature contribution compared to the other two categories (Figure 8C).

We performed ten rounds of predictions on HL-negative genes not included in the corresponding training datasets using the models, identifying 222 genes consistently predicted as positive in all instances (Figure 8D). Among these, 18 genes exhibited HL phenotypes in mouse models (Figure 8E). Further analysis of patient data from the CDGC cohort and experimental validations pinpointed *ERCC2* and *TBX2* as novel HL genes ^54^.

## Discussion

By integrating extensive data from numerous public repositories and the HL patients from the CDGC cohort, GDC substantially advanced the understanding of the genetic landscape of hearing loss. Analyses of the GDC dataset refined the evidence for interpreting variant pathogenicity in specific genes and identified six novel mutational hotspots within key HL genes, highlighting the dynamic nature of genomic regions linked to auditory dysfunction. These insights significantly enrich our diagnostic toolkit, enabling the precise identification of pathogenic variants that can be directly implemented in clinical settings to improve both diagnostic accuracy and patient outcomes. Moreover, further exploration of the GDC dataset through comparative phenotypic analysis highlighted significant disparities between human and mouse models. Detailed examination of gene expression in the cochleae of both mice and rhesus macaques illuminated the conservation and divergence of genes essential for auditory function and development. Leveraging in-depth data on gene expression, function, and phenotypes, the application of the machine learning approach identified 18 candidate HL genes. Notably, among these candidates, *TBX2* and *ERCC2*, have been validated as novel HL genes using the patient data from the CDGC cohort, representing a landmark advancement in understanding HL genetics ^54^.

The rapid advancement of DNA sequencing technologies has markedly improved our ability to pinpoint genes and mutations associated with diseases. However, this progress also presents a significant challenge in accurately interpreting the pathogenicity of numerous rare variants. This issue is also evident in the GDC, where up to 95.2% of variants are either of uncertain significance or remain unclassified, highlighting the need for comprehensive integration of clinical, genetic, and molecular functional data to bridge these gaps. In response, the American College of Medical Genetics and Genomics (ACMG) and the Association for Molecular Pathology (AMP) jointly developed guidelines to standardize genetic diagnostic procedures, incorporating up to 28 lines of evidence to facilitate variant interpretation and classification ^55^. This initiative has markedly enhanced the consistency, reproducibility, and transparency of genetic diagnosis. Adapting these guidelines to accommodate the extensive diversity across diseases and genes remains critical for meeting the intricate demands of clinical practice and research. To this end, specific evidence rules, such as PVS1 ^56^, PP3 ^57^, BP4 ^58^, and PS4 ^59^, along with specific guidelines for particular disorders like inherited cardiomyopathy ^60^, cerebral creatine deficiency syndromes ^61^, and genetic HL ^62^, have been continuously optimized and refined over recent years. Presently, the GDC covers 17 out of 24 lines of evidence applicable to genetic HL as outlined by HL-ACMG ^62^, providing substantial support for clinical genetic diagnosis.

More importantly, the GDC provided new insights that enhance the application of the ACMG/AMP guidelines for clinical genetic diagnosis. The PM1 rule, defined as the significant enrichment of pathogenic missense variants in a mutational hotspot and/or critical and well-established functional domain, represents an important line of pathogenic evidence. However, the practical implementation of the PM1 was constrained by a scarcity of studies on the identification of such hotspots. In the context of genetic HL, only collagen genes and the *KCNQ4* gene were previously reported to have mutational hotspots amenable to PM1 ^50,62^. In this study, we systematically mapped the distribution of pathogenic missense variants at the amino acid level across 201 HL genes, identifying 12 genes with hotspots enriched for pathogenic variants. According to Tavtigian et al ^49^, the positive likelihood ratio of eight enriched genes, including six reported for the first time in this study, exceed thresholds of supporting or moderate strength level, thus broadening the applicability of this rule. Notably, among all 12 transcription factor genes examined, 10 exhibited enrichments in DNA binding domains, highlighting the critical amino acids within these domains that impact transcriptional efficacy. By pinpointing these hotspots, this discovery provides valuable insights into the genetic landscape of deafness and establishes a foundation for subsequent studies on the functional implications of HL genes. Another specific rule addressed by this study focused on NMD-escape premature termination codon variants, which present challenges in genetic diagnosis due to their varied impacts on protein function. A key determinant of NMD efficiency is the position of PTVs, encapsulated by the 50 nucleotide (50nt) boundary rule: variants occurring more than 50 to 55 nucleotides upstream of the last exon junction are targeted for degradation ^63,64^. Conversely, PTVs located downstream of this boundary are considered NMD-escape and are subject to a reduction in the strength of the pathogenic PVS1 rule for genetic diagnostics ^56^. The pathogenic mechanisms of NMD-escape PTVs may include the production of a truncated protein leading to a LoF effect, or alternatively, the truncated protein may exert a dominant negative or a GoF effect, with impacts varying according to the specific gene and variant ^65^. This study identified a significant number of predicted NMD-escape PTV variants in multiple HL genes, including *MARVELD2*, *MYH9*, and *MYH14*. The results suggest that subsequent identification of NMD-escape PTVs in these genes is not subject to reduction of evidence strength, it also revealed the function implication that the C-terminal amino acids of these proteins are essential for their functionality, and disruption at these sites may compromise the entire protein function.

Constructing animal models that accurately simulate human phenotypes is essential for validating disease genes and exploring underlying mechanisms. The mouse model has been widely used in hearing research due to its short generation time and the structural similarity of its cochlea to that of humans, which allows for effective dissection and imaging ^66^. Despite this, discrepancies between human and mouse phenotypes are not uncommon. For instance, GDC data on mouse models for 201 human HL genes revealed that up to 40.3% of these genes did not associate with the expected deaf phenotype in mice. Interestingly, these discrepancies appear minimally related to the sequence homology of the corresponding proteins or gene expression levels, suggesting that they may stem from a higher order of genomic evolution and organization ^51^. Comparative genomics and transcriptomics studies have indicated that the phenotypic differences between species are partly due to variations in non-coding regulatory elements and transcriptome profiles, rather than differences in protein-coding sequences ^52,67^. Furthermore, the genetic background of different mouse strains significantly affects phenotypic expression, as exemplified by the *Atp6v1b1* gene, which failed to induce HL when knocked out in the C57 background but succeeded in the MRL/MpJ background ^68^. Genetic redundancy, where a gene deletion or mutation is compensated for by other genes, may also influence phenotypic variations ^69^. To overcome these challenges, strategies such as constructing humanized mouse models, knocking out additional genes within the same family, selecting different mouse strains, and incorporating environmental factors have been proposed. Successful humanization of mouse models for deafness genes, like *MYO6* ^70^ and *COCH* ^71^, has yielded critical insights into the molecular mechanisms of hearing diseases.

Research shows that a significant proportion of orthologous protein-coding genes maintain conserved expression patterns between humans and mice ^67^, a result that aligns with our findings of strong correlation in gene expression levels between mouse and rhesus macaque cochlear tissues. Leveraging these insights, we integrated gene expression data from both species, incorporating expression correlations, pathways, and functional data into the feature set. Using the SMOTE-Random Forest Classifier to manage oversampled datasets, we developed a scoring model to identify candidate genes for HL. The model was primarily driven by gene expression levels across different cell types, indicating potential improvements by including more types of omics data such as proteomics and epigenomics ^72,73^. Subsequent overlapping of HL phenotype data from mouse models facilitated the filtering of candidate genes. Through genetic analyses within the CDGC cohort and subsequent experimental validation, we confirmed two novel HL genes, *TBX2* and *ERCC2*^54^. These results underscored the value of extensive, multi-layered genetic and genomic data in the GDC, demonstrating how the use of such data can aid in uncovering novel disease genes and deepen our understanding of disease mechanisms.

The results of this study should be considered in light of several limitations. A significant proportion of variants within the GDC remain unclassified or of uncertain significance, which could potentially obscure our understanding of the genetic landscape of HL. Moreover, the GDC exhibited only a limited number of pathogenic non-coding variants across a handful of genes. The analysis of non-coding variants is notably challenging due to their intricate regulatory roles and the absence of clear clinical correlations. To improve the accuracy of interpreting both coding and non-coding variants, it is crucial to integrate more comprehensive genetic data from a broad range of ethnic groups and to innovate in the development of new high-throughput functional assays and analytic algorithms. Another critical obstacle is the difficulty in obtaining inner ear tissue samples, which restricts the generation of comprehensive epigenetic and expression data that are essential for a thorough understanding of the molecular mechanisms underlying auditory functions and deafness. Addressing these limitations is crucial for future research efforts aimed at advancing our capabilities to explore and interpret auditory-related issues and developing effective therapeutic strategies for deafness.

In conclusion, GDC represents a valuable endeavor, serving as a unified and enriched information repository that comprehensively captures data and knowledge to specifically advance research and clinical practices in audiology and hearing-related conditions. Future enhancements of the GDC will focus on expanding coverage to additional HL-related genes, incorporating epigenetic regulatory information of cochlea, and developing the knowledge graph through literature mining, all of which will further solidify its role as an invaluable resource for the research community.

## Material and Methods

### CDGC Cohort Study

The CDGC project is a China-wide cooperative network of medical genetics research on hearing loss and related disorders. The aim of the study is to elucidate the role of genetic variation in the pathogenesis of HL by bringing together the increasing genetic information and high-quality clinical data, as well as analysis expertise, to discover the basis for the prevention and treatment of HL, and to exploit the new knowledge to reduce the burden of disease. Since 2013, the CDGC project has involved 22,125 cases and 7,254 controls across mainland China. The exclusion criteria were: 1) conductive HL (e.g., HL secondary to otitis media, chronic myringitis, perforated eardrum, and tympanosclerosis); 2) presbycusis (age of onset > 40 years); 3) unilateral HL without a family history; and 4) mild HL without a family history. The involved cases went through three-stage genetic testing using SNPscan, CDGC-HL panel, and whole genome sequencing (WGS) ^74^. Variant interpretation and genetic diagnosis were performed on up to 201 HL genes based on the set of ACMG/AMP guidelines and the updates ^55–57,62,75,76^, in the context of patient phenotype, onset age, HL severity, other otologic testing records, family history, and medication history. Putative diagnostic variants were Sanger sequenced for validation. All variants identified in the CDGC cohort in 201 HL genes were included in the GDC, with information on pathogenicity classifications and allele frequencies.

### Data Integration and Development of GDC

GDC integrates genetic and phenotypic information from 51 public databases. Each gene was annotated with 11 aspects of information from 29 databases, including summary information, phenotype and disease correlation, gene expression, subcellular localization, protein-protein interaction, gene function and pathway, animal model, as well as gene-drug interaction. Variants in GDC were incorporated from multiple data sources in addition to CDGC, including ClinVar, DVD, HGMD, gnomAD, and ChinaMAP. A total of 27 databases provides variant annotations, including functional consequence and impact, allele frequencies in different populations, pathogenicity classification, computational prediction scores, correlated diseases or phenotypes, and linked publications. Variant Effect Predictor (VEP) was employed to provide comprehensive annotations for each variant. All data were standardized and stored in the MongoDB database. The GDC website (http://gdcdb.net/) was developed using the Python-Flask, Nginx, and React JavaScript frameworks, with UI and components built using Antd Design. The web interface of GDC provides user-friendly searching functions that were equipped to automatically recognize four types of key terms: gene symbols, genomic coordinates of variants, phenotype terms, and disease names. In addition, GDC offers downloadable datasets in VCF/BED/GFF formats for offline annotation.

### Pathogenic Missense Variant Density Estimation and Likelihood Ratio Calculation

P/LP missense variants from CDGC, ClinVar, DVD, and HGMD, and non-P/LP missense variants from the gnomAD population with maximum MAF < 0.0001 were respectively utilized to estimate the distribution density in terms of the amino acid location. Gaussian kernel density from the NumPy package was used to estimate the density of missense variants and to draw the density map for each gene. The enrichment region was determined based on the location of missense variants and protein function domains retrieved from InterPro. The bootLR ^77^ and epiR ^78^ packages were used to calculate the likelihood ratio (LR), odds ratio (OR), and Fisher’s exact p-value of enrichment for each region, employing the following formulas:

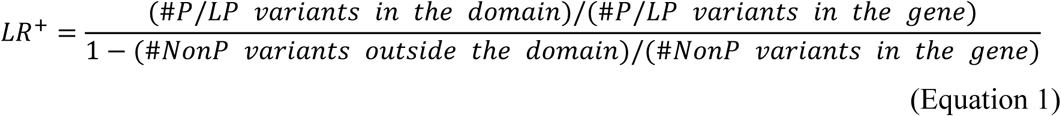

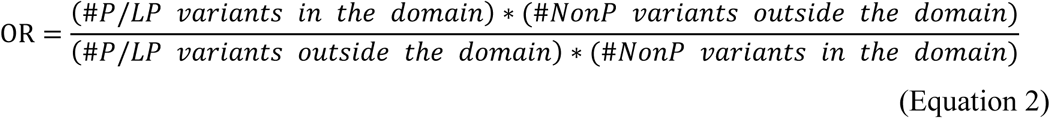

where *#* is the number of variants and NonP indicates non-pathogenic variants.

### Single-Cell Expression Analysis of the Mouse Cochlea

Single-cell transcriptomic data of mouse cochlea was retrieved from GEO (Accession ID: GSE172110, GSE182202, GSE181454, and GSE202920; GSA: CRA004814). We analyzed the expression profiles of single cells from the mouse cochlea at different developmental stages: E14, E18, P1, P8, P14, P20, P28, M1, M2, M5, M12, and M15. The initial normalization and clustering were performed by Seurat v4.2.1^79^. The matrix of read-count data of each cell for each gene was loaded individually and converted to Seurat objects. Harmony package was used to integrate multiple datasets ^80^. Briefly, gene expression matrices were loaded as count matrices, and a pooled Seurat object was generated on the specified count matrices. Filtering, normalization, and scaling were performed, and integration was conducted by the ‘RunHarmony’ function which iteratively integrated cells to the point of convergence. For integrated datasets, harmony embeddings were used for all downstream analyses. The integrated data set was then used for downstream dimensionality reduction and clustering analyses. Total cell clustering was performed by the “FindClusters” function to define cell identity. Dimensionality reduction was performed with the “RunUMAP” function. Marker genes for each cluster were determined with the Wilcoxon rank-sum test by the “FindAllMarkers” function. Only those with “avg_logFC” > 0.5 and “p_val_adj” < 0.05 were considered as marker genes. Cell types were identified based on the expression of classic marker genes.

### RNA-Seq of Rhesus Macaque Cochlea and Data Analysis

Three cochlear samples were retrieved from two rhesus macaques via dissection. Total RNA was extracted from these samples, and mRNA was subsequently enriched using Oligo(dT) magnetic beads. A fragmentation buffer was added to break the mRNA into short fragments. These fragments served as templates for synthesizing the first strand of cDNA using random hexamer primers. Subsequent steps involved adding buffer, dNTPs, RNase H, and DNA polymerase I to synthesize the second cDNA strand. The cDNA was then purified using the QIAquick PCR Purification Kit (Qiagen) and eluted with EB buffer. End repair processes were followed by the addition of poly(A) tails and ligation of sequencing adapters. The desired fragment sizes were selected via agarose gel electrophoresis, followed by PCR amplification. The prepared sequencing library was then sequenced on the Illumina HiSeq platform.

Raw reads in FASTQ file format were filtered for quality control using fastp and FastQC. After confirming the quality, clean reads were mapped to the Macaca mulatta genome using Hisat2. The mapped reads were quantified using StringTie. Transcripts per million (TPM) normalized the RNA-Seq data, calculated as the read number of a transcript divided by the total clone count of the sample, then multiplied by 10^6^. Differential gene expression between the samples was determined using the EdgeR.

### Candidate HL Gene Prediction using SMOTE-Random Forest Classifier

To identify candidate HL genes, the machine learning approach was utilized on data integrated by GDC. The initial dataset comprised genes characterized by 275 features, including expression levels in mouse and rhesus macaque cochlea, expression correlation with confirmed HL genes, GO terms, and pathways. Confirmed HL genes were labeled positive, while those not listed in the HPO as having HL as a minor symptom were labeled negative. The negative genes were randomly divided into ten subsets, each combined with the positive genes to form 10 training datasets. Given the imbalance in the ratio of positive to negative labels (approximately 9:1) in each dataset, the Synthetic Minority Over-sampling Technique (SMOTE) was applied to oversample the positive class. A Random Forest Classifier was employed to generate 10 models, each predicting the labels of genes not included in its training dataset, yielding nine prediction outcomes for each negative gene. To synthesize these results, the predictions from the ten models were aggregated, counting the number of times each gene was predicted as positive across all models (ranging from 0 to 9 times). This count served as a score indicating the likelihood of a gene being associated with HL, with higher scores suggesting a greater probability. Furthermore, a mean decrease impurity analysis was conducted for the 275 features in each model, yielding an importance score for each feature. A higher feature score signified a more significant influence on the model, indicating a pivotal role in the classification task.

## Acknowledgments

None.

## Funding

This research was financially supported by the National Natural Science Foundation of China (82171836), the Science and Technology Department of Sichuan Province (2024NSFSC0648) and the 1·3·5 Project for Disciplines of Excellence, West China Hospital, Sichuan University (ZYJC20002).

## Author contributions

**Hui Cheng** Investigation, Formal analysis, Data collection and standardization, Writing - Original draft. **Mingjun Zhong:** Formal analysis, Data collection and standardization, Writing - Review & Editing. **Jia Geng:** Formal analysis, Data collection and standardization, Writing - Review & Editing. **Xuegang Wang**: Website development, Website optimization, Website testing. **Wenjian Li** and **Kanglu Pei:** Formal analysis, Writing - Review & Editing. **Yu Lu**, **Jing Cheng**, **Fengxiao Bu** and **Huijun Yuan**: Study design, Writing - Review & Editing, Supervision, Project administration, Funding acquisition.

## Competing interests

The authors declare no competing financial interests.

## Notes

### Competing Interest Statement

The authors have declared no competing interest.

## References

1. Smith, R. J., Bale, J. F. & White, K. R. Sensorineural hearing loss in children. The Lancet 365, 879–890 (2005).

2. Korver, A. M. H. et al. Congenital hearing loss. Nat Rev Dis Primers 3, 16094 (2017).

3. Lieu, J. E. C., Kenna, M., Anne, S. & Davidson, L. Hearing Loss in Children: A Review. JAMA 324, 2195–2205 (2020).

4. Gana, N. et al. Congenital Cytomegalovirus-Related Hearing Loss. Audiol Res 14, 507–517 (2024).

5. Azaiez, H. et al. Genomic Landscape and Mutational Signatures of Deafness-Associated Genes. The American Journal of Human Genetics 103, 484–497 (2018).

6. Huang, S. et al. Gene4HL: An Integrated Genetic Database for Hearing Loss. Front Genet 12, 773009 (2021).

7. Van, C. G. & Smith, R. Hereditary Hearing Loss Homepage. Hereditary Hearing Loss Homepage https://hereditaryhearingloss.org (2022).

8. Patel, M. J. et al. Disease-specific ACMG/AMP guidelines improve sequence variant interpretation for hearing loss. Genet Med 23, 2208–2212 (2021).

9. Kim, S. Y. et al. Improving genetic diagnosis by disease-specific, ACMG/AMP variant interpretation guidelines for hearing loss. Sci Rep 12, 12457 (2022).

10. Kremer, H. Novel gene discovery for hearing loss and other routes to increased diagnostic rates. Hum Genet 141, 383–386 (2022).

11. Petit, C., Bonnet, C. & Safieddine, S. Deafness: from genetic architecture to gene therapy. Nat Rev Genet 24, 665–686 (2023).

12. Kolla, L. et al. Characterization of the development of the mouse cochlear epithelium at the single cell level. Nat Commun 11, 2389 (2020).

13. Raney, B. J. et al. The UCSC Genome Browser database: 2024 update. Nucleic Acids Res 52, D1082–D1088 (2024).

14. Martin, F. J. et al. Ensembl 2023. Nucleic Acids Res 51, D933–D941 (2023).

15. Orvis, J. et al. gEAR: Gene Expression Analysis Resource portal for community-driven, multi-omic data exploration. Nat Methods 18, 843–844 (2021).

16. GTEx Consortium. Genetic effects on gene expression across human tissues. Nature 550, 204–213 (2017).

17. UniProt Consortium. UniProt: the Universal Protein Knowledgebase in 2023. Nucleic Acids Res 51, D523–D531 (2023).

18. Gene Ontology Consortium et al. The Gene Ontology knowledgebase in 2023. Genetics 224, iyad031 (2023).

19. Paysan-Lafosse, T. et al. InterPro in 2022. Nucleic Acids Res 51, D418–D427 (2023).

20. Burley, S. K. et al. Protein Data Bank (PDB): The Single Global Macromolecular Structure Archive. Methods Mol Biol 1607, 627–641 (2017).

21. Milacic, M. et al. The Reactome Pathway Knowledgebase 2024. Nucleic Acids Res 52, D672–D678 (2024).

22. Kanehisa, M., Furumichi, M., Tanabe, M., Sato, Y. & Morishima, K. KEGG: new perspectives on genomes, pathways, diseases and drugs. Nucleic Acids Res 45, D353–D361 (2017).

23. Szklarczyk, D. et al. The STRING database in 2023: protein-protein association networks and functional enrichment analyses for any sequenced genome of interest. Nucleic Acids Res 51, D638–D646 (2023).

24. Oughtred, R. et al. The BioGRID database: A comprehensive biomedical resource of curated protein, genetic, and chemical interactions. Protein Sci 30, 187–200 (2021).

25. Gong, L., Whirl-Carrillo, M. & Klein, T. E. PharmGKB, an Integrated Resource of Pharmacogenomic Knowledge. Curr Protoc 1, e226 (2021).

26. Cannon, M. et al. DGIdb 5.0: rebuilding the drug-gene interaction database for precision medicine and drug discovery platforms. Nucleic Acids Res 52, D1227–D1235 (2024).

27. Karczewski, K. J. et al. The mutational constraint spectrum quantified from variation in 141,456 humans. Nature 581, 434–443 (2020).

28. Cao, Y. et al. The ChinaMAP analytics of deep whole genome sequences in 10,588 individuals. Cell Res 30, 717–731 (2020).

29. Kubo, M. & Guest Editors. BioBank Japan project: Epidemiological study. J Epidemiol 27, S1 (2017).

30. Landrum, M. J. et al. ClinVar: public archive of relationships among sequence variation and human phenotype. Nucleic Acids Res 42, D980–985 (2014).

31. Stenson, P. D. et al. Human Gene Mutation Database: towards a comprehensive central mutation database. J Med Genet 45, 124–126 (2008).

32. Rentzsch, P., Witten, D., Cooper, G. M., Shendure, J. & Kircher, M. CADD: predicting the deleteriousness of variants throughout the human genome. Nucleic Acids Res 47, D886–D894 (2019).

33. Ioannidis, N. M. et al. REVEL: An Ensemble Method for Predicting the Pathogenicity of Rare Missense Variants. Am J Hum Genet 99, 877–885 (2016).

34. Jaganathan, K. et al. Predicting Splicing from Primary Sequence with Deep Learning. Cell 176, 535–548.e24 (2019).

35. Allot, A. et al. Tracking genetic variants in the biomedical literature using LitVar 2.0. Nat Genet 55, 901–903 (2023).

36. Lin, Y.-H. et al. variant2literature: full text literature search for genetic variants. Preprint at 10.1101/583450 (2019).

37. Gargano, M. A. et al. The Human Phenotype Ontology in 2024: phenotypes around the world. Nucleic Acids Res 52, D1333–D1346 (2024).

38. Miller, N., Lacroix, E. M. & Backus, J. E. MEDLINEplus: building and maintaining the National Library of Medicine’s consumer health Web service. Bull Med Libr Assoc 88, 11–17 (2000).

39. Amberger, J. S. & Hamosh, A. Searching Online Mendelian Inheritance in Man (OMIM): A Knowledgebase of Human Genes and Genetic Phenotypes. Curr Protoc Bioinformatics 58, 1.2.1–1.2.12 (2017).

40. Rehm, H. L. et al. ClinGen--the Clinical Genome Resource. N Engl J Med 372, 2235–2242 (2015).

41. Eppig, J. T. Mouse Genome Informatics (MGI) Resource: Genetic, Genomic, and Biological Knowledgebase for the Laboratory Mouse. ILAR J 58, 17–41 (2017).

42. Tasdemir-Yilmaz, O. E. et al. Diversity of developing peripheral glia revealed by single-cell RNA sequencing. Dev Cell 56, 2516–2535.e8 (2021).

43. Iyer, A. A. et al. Cellular reprogramming with ATOH1, GFI1, and POU4F3 implicate epigenetic changes and cell-cell signaling as obstacles to hair cell regeneration in mature mammals. Elife 11, e79712 (2022).

44. Xue, N. et al. Genes related to SNPs identified by Genome-wide association studies of age-related hearing loss show restriction to specific cell types in the adult mouse cochlea. Hear Res 410, 108347 (2021).

45. Xu, Z. et al. Profiling mouse cochlear cell maturation using 10×Genomics single-cell transcriptomics. Front Cell Neurosci 16, 962106 (2022).

46. Sun, G. et al. Single-cell transcriptomic atlas of mouse cochlear aging. Protein Cell 14, 180–201 (2023).

47. Rivas, M. A. et al. Human genomics. Effect of predicted protein-truncating genetic variants on the human transcriptome. Science 348, 666–669 (2015).

48. Cheng, J. et al. Functional mutation of SMAC/DIABLO, encoding a mitochondrial proapoptotic protein, causes human progressive hearing loss DFNA64. Am J Hum Genet 89, 56–66 (2011).

49. Tavtigian, S. V. et al. Modeling the ACMG/AMP variant classification guidelines as a Bayesian classification framework. Genet Med 20, 1054–1060 (2018).

50. Naito, T. et al. Comprehensive genetic screening of KCNQ4 in a large autosomal dominant nonsyndromic hearing loss cohort: genotype-phenotype correlations and a founder mutation. PLoS One 8, e63231 (2013).

51. Cardoso-Moreira, M. et al. Developmental Gene Expression Differences between Humans and Mammalian Models. Cell Rep 33, 108308 (2020).

52. Han, S. K., Kim, D., Lee, H., Kim, I. & Kim, S. Divergence of Noncoding Regulatory Elements Explains Gene-Phenotype Differences between Human and Mouse Orthologous Genes. Mol Biol Evol 35, 1653–1667 (2018).

53. Liao, B.-Y. & Zhang, J. Null mutations in human and mouse orthologs frequently result in different phenotypes. Proc Natl Acad Sci U S A 105, 6987–6992 (2008).

54. Hua, W. et al. Frameshift variants in TBX2 underlie autosomal-dominant hearing loss with incomplete penetrance of nystagmus. Preprint at 10.1101/2024.07.18.24310488 (2024).

55. Richards, S. et al. Standards and guidelines for the interpretation of sequence variants: a joint consensus recommendation of the American College of Medical Genetics and Genomics and the Association for Molecular Pathology. Genet Med 17, 405–424 (2015).

56. Abou Tayoun, A. N., et al. Recommendations for interpreting the loss of function PVS1 ACMG/AMP variant criterion. Hum Mutat 39, 1517–1524 (2018).

57. Pejaver, V. et al. Calibration of computational tools for missense variant pathogenicity classification and ClinGen recommendations for PP3/BP4 criteria. Am J Hum Genet 109, 2163–2177 (2022).

58. Stenton, S. L. et al. Assessment of the evidence yield for the calibrated PP3/BP4 computational recommendations. Genet Med 101213 (2024) doi:10.1016/j.gim.2024.101213.

59. Liu, S. et al. Quantitative thresholds for variant enrichment in 13,845 cases: improving pathogenicity classification in genetic hearing loss. Genome Med 15, 116 (2023).

60. Kelly, M. A. et al. Adaptation and validation of the ACMG/AMP variant classification framework for MYH7-associated inherited cardiomyopathies: recommendations by ClinGen’s Inherited Cardiomyopathy Expert Panel. Genetics in Medicine 20, 351–359 (2018).

61. Goldstein, J. et al. ClinGen variant curation expert panel recommendations for classification of variants in GAMT, GATM and SLC6A8 for cerebral creatine deficiency syndromes. Mol Genet Metab 142, 108362 (2024).

62. Oza, A. M. et al. Expert specification of the ACMG/AMP variant interpretation guidelines for genetic hearing loss. Hum Mutat 39, 1593–1613 (2018).

63. Chang, Y.-F., Imam, J. S. & Wilkinson, M. F. The nonsense-mediated decay RNA surveillance pathway. Annu Rev Biochem 76, 51–74 (2007).

64. Lewis, B. P., Green, R. E. & Brenner, S. E. Evidence for the widespread coupling of alternative splicing and nonsense-mediated mRNA decay in humans. Proc Natl Acad Sci U S A 100, 189–192 (2003).

65. Torene, R. I. et al. Systematic analysis of variants escaping nonsense-mediated decay uncovers candidate Mendelian diseases. Am J Hum Genet 111, 70–81 (2024).

66. Carlson, R. J. & Avraham, K. B. Emerging complexities of the mouse as a model for human hearing loss. Proc Natl Acad Sci U S A 119, e2211351119 (2022).

67. Breschi, A., Gingeras, T. R. & Guigó, R. Comparative transcriptomics in human and mouse. Nat Rev Genet 18, 425–440 (2017).

68. Tian, C. et al. Hearing loss without overt metabolic acidosis in ATP6V1B1 deficient MRL mice, a new genetic model for non-syndromic deafness with enlarged vestibular aqueducts. Hum Mol Genet 26, 3722–3735 (2017).

69. Peng, J. Gene redundancy and gene compensation: An updated view. J Genet Genomics 46, 329–333 (2019).

70. Wang, J. et al. A humanized mouse model, demonstrating progressive hearing loss caused by MYO6 p.C442Y, is inherited in a semi-dominant pattern. Hear Res 379, 79–88 (2019).

71. Verdoodt, D. et al. Rational design of a genomically humanized mouse model for dominantly inherited hearing loss, DFNA9. Hear Res 442, 108947 (2024).

72. Chen, Y., Wu, X. & Jiang, R. Integrating human omics data to prioritize candidate genes. BMC Med Genomics 6, 57 (2013).

73. Chung, J. et al. Genome-wide association and multi-omics studies identify MGMT as a novel risk gene for Alzheimer’s disease among women. Alzheimers Dement (2022) doi:10.1002/alz.12719.

74. Du, W. et al. A rapid method for simultaneous multi-gene mutation screening in children with nonsyndromic hearing loss. Genomics 104, 264–270 (2014).

75. Biesecker, L. G. et al. ClinGen guidance for use of the PP1/BS4 co-segregation and PP4 phenotype specificity criteria for sequence variant pathogenicity classification. Am J Hum Genet 111, 24–38 (2024).

76. Walker, L. C. et al. Using the ACMG/AMP framework to capture evidence related to predicted and observed impact on splicing: Recommendations from the ClinGen SVI Splicing Subgroup. Am J Hum Genet 110, 1046–1067 (2023).

77. Marill, K. A., Chang, Y., Wong, K. F. & Friedman, A. B. Estimating negative likelihood ratio confidence when test sensitivity is 100%: A bootstrapping approach. Stat Methods Med Res 26, 1936–1948 (2017).

78. Stevenson, M. & Sergeant, E. epiR: Tools for the Analysis of Epidemiological Data. 2.0.75 10.32614/CRAN.package.epiR (2008).

79. Hao, Y. et al. Integrated analysis of multimodal single-cell data. Cell 184, 3573–3587.e29 (2021).

80. Korsunsky, I. et al. Fast, sensitive and accurate integration of single-cell data with Harmony. Nat Methods 16, 1289–1296 (2019).

